# Neuronal fatty acid oxidation fuels memory after intensive learning

**DOI:** 10.1101/2024.12.18.629183

**Authors:** Alice Pavlowsky, Bryon Silva, Lydia Danglot, Pierre-Yves Plaçais, Thomas Preat

**Affiliations:** Energy & Memory, Brain Plasticity Unit, CNRS, ESPCI Paris, PSL Research University; Paris, France; Université Paris Cité, NeurImag Imaging Facility, Institute of Psychiatry and Neuroscience of Paris, INSERM U1266; Paris, France

## Abstract

Metabolic flexibility allows cells to adapt to different fuel sources, which is particularly important for cells with high metabolic demands. In contrast, neurons, which are major energy consumers, are considered to rely almost solely on glucose and its derivatives to support their metabolism^1–3^. Here, using *Drosophila melanogaster*, we show memory formed after intensive massed training is dependent on mitochondrial fatty acid (FA) β-oxidation to produce ATP in neurons of the mushroom bodies (MB), a major integrative center in insects’ brain. We identify neuronal lipid droplets as the main source of FAs for this type of memory. Furthermore, we demonstrate that this intensive massed training is associated with mitochondria network remodeling in the soma of MB neurons, resulting in increased mitochondrial size. Artificially increasing mitochondria size in adult MB neurons increases ATP production in their soma and, at the behavioral level, strikingly results in improved memory performance after massed training. These findings challenge the prevailing view that neurons are unable to use FAs for energy production, and importantly revealing on the contrary that *in vivo* neuronal FA oxidation has an essential role in cognitive function, including memory formation.

## Main Text

In mammals, FA β-oxidation contributes to up to 20% of total brain oxidative metabolism^4^, and glial cells are viewed as the major brain cell type to use FAs for energy production^5–8^. On the other hand, FAs are considered to be metabolic substrates that promote mitochondrial reactive oxygen species (ROS) generation, to which neurons are especially sensitive^6,7^, an argument that has been put forward to support the predominant view that neurons do not perform β-oxidation^9^. Despite the toxicity associated with FA metabolism, there is increasing evidence of an FA internal store in neurons^10^, and it was recently suggested *in vitro* that impairing lipolysis from this internal store impairs neuronal energy production^11,12^. However, whether neuronal mitochondria actually perform β-oxidation *in vivo* remains unknown.

To address this question, we used classical pavlovian olfactory conditioning in *Drosophila melanogaster.* In this olfactory memory paradigm, repeated consecutive learning sessions (intensive massed training) induce a memory that decays within 1-2 days, while repeated sessions spaced in time (intensive spaced training) lead to a memory that lasts for up to one week^13^. This latter type of memory, classically called long-term memory (LTM), is dependent on protein synthesis^13^ and requires an extended post-training upregulation of pyruvate mitochondrial metabolism in MB neurons^14,15^, the major associative memory center in Drosophila^16^. In contrast, the memory formed after intensive massed training, which resembles cramming-like learning, is classically called anesthesia-resistant memory (ARM)^13^ and is not dependent on mitochondrial pyruvate metabolism^14^. Furthermore, this memory formed after massed training does not depend on ketone body oxidation in fed flies, an alternative fuel used by MB neurons for memory formation under starved conditions^17^. We therefore hypothesized that to support memory formed after massed training, MB neurons would use FAs as an energy substrate. To test this hypothesis, we first targeted the system that imports FAs into the mitochondria. Long-chain FAs must first be activated to acyl-coenzyme A (Acyl-CoA) esters by cytosolic acyl-CoA synthetase before they can be directed into the mitochondria by the carnitine shuttle system^18,19^ (Fig. 1a). Since the outer mitochondrial membrane component of this shuttle, carnitine palmitoyltransferase 1 (CPT1), catalyzes the rate-limiting step of FA import^20^, we tested its involvement in memory using an RNAi-based knock-down (KD). To restrict the expression of CPT1 RNAi to adult MB neurons, we used the VT30559-Gal4 driver^14^ in combination with the ubiquitously expressed thermosensitive Gal4 inhibitor, Gal80 (Tub-Gal80^ts^)^21^. Placing the flies at 30°C for 2-3 days therefore allows RNAi expression in MB neurons (see Methods for details). Flies were then subjected to an intensive massed aversive training and tested for memory performance 24 h later. Downregulation of CPT1 expression in adult MB impaired memory after massed training, whereas 24-h memory formed after spaced training was normal (Fig. 1b). Memory after massed training was normal in a control experiment in which CPT1 RNAi expression was not induced (Extended Data Fig. 1a). Finally, CPT1 KD did not alter innate shock reactivity or olfactory acuity (Supplementary Table 1). These results were replicated with a second non-overlapping RNAi line targeting CPT1 (Extended Data Fig. 1b, Supplementary Table 1). For simplicity, the combined phenotypes described above are hereafter described as a specific defect in memory after massed training. Both CPT1 RNAi lines, previously validated in glial cells^17^, efficiently reduced the CPT1 mRNA level in Drosophila brains when expressed specifically in neurons (Supplementary Table 2). These results show that FAs imported into the mitochondria via the carnitine shuttle are required specifically for memory formed after massed training.

**Figure 1:**
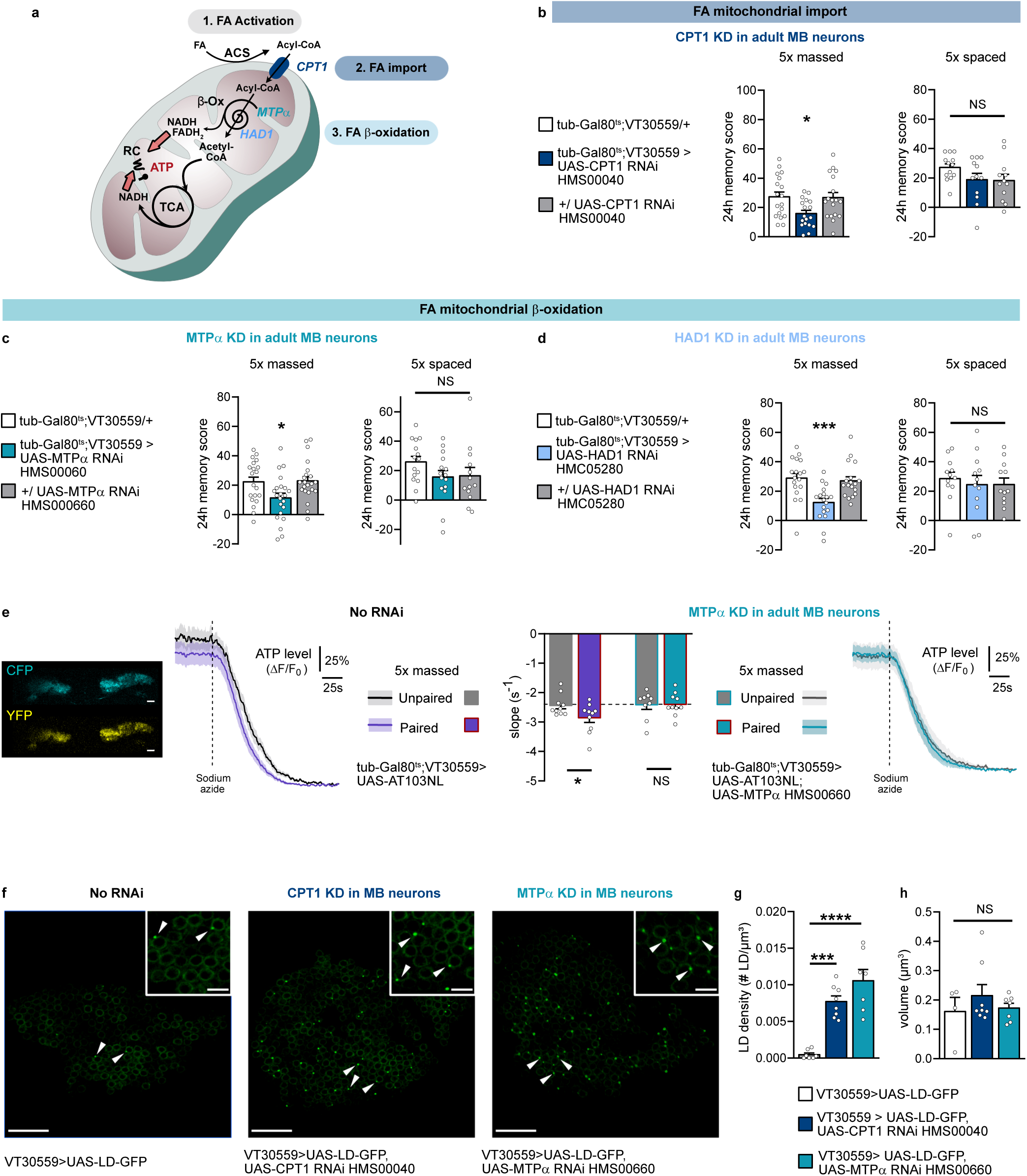
FA mitochondrial β-oxidation is required in MB neurons for memory after massed training. **a.** Prior to mitochondrial import, FAs are activated by acyl-CoA synthetase (ACS) into acyl-CoA, which is then shuttled into the mitochondrial matrix by the carnitine shuttle system. The outer mitochondrial membrane component of this shuttle, carnitine palmitoyltransferase 1 (CPT1), catalyzes the rate-limiting step of FA import. β-oxidation (β-ox) is a cyclic process: each cycle shortens the acyl-CoA by two carbons and releases an acetyl-CoA together with reduced cofactors (FADH_2_ and NADH) that directly feed into (red arrows) the respiratory chain (RC) to produce ATP. The β-oxidation machinery harbors different chain length–specific enzymes organized into different functional complexes: MTP1 is part of the mitochondrial trifunctional protein (MTP), an enzymatic complex attached to the inner mitochondrial membrane that shortens long-chain FAs to medium-chain lengths; they are then oxidized by a matrix system containing soluble enzymes for medium- and short-chain FAs including HAD1, which shares homology with the mammalian 3-hydroxyacyl-CoA dehydrogenases. Acetyl-CoA can be further oxidized within the tricarboxylic acid cycle (TCA) to produce reduced cofactors that fuel the RC for ATP production. **b.** Inhibition of FA mitochondrial import by CPT1 (CPT1 KD) in adult MB neurons impaired memory after massed training (n=18, F_2,51_=4.40, P<0.05), but not after spaced training (n=12-13, F_2,34_=1.86, P=0.17). **c-d.** Inhibition of FA mitochondrial β-oxidation in adult MB neurons by either Mtpα (c) or HAD1 (d) KD impaired memory after massed training (b: n=20-22, F_2,61_=4.43, P<0.05, c: n=18, F_2,51_=9.43, P<0.001), but not after spaced training (b: n=13-15, F_2,40_=1.56, P=0.22, c: n=12, F_2,33_=0.21, P=0.81). **e.** The ATP FRET sensor AT103.NL was expressed in adult MB neurons, visualized in the CFP and YFP channels and quantified in the soma region of MB neurons (scale bar: 15 μm). Application of 5 mM of sodium azide (dashed line) resulted in a fast decrease in the FRET ratio (mean trace ± s.e.m.), making it possible to estimate the level of ATP consumption before sodium azide application by quantifying the slope of the FRET ratio decrease. After massed associative training (purple), the rate of the resulting ATP decrease was faster than in flies subjected to the non-associative protocol (grey; n=10, t_18_=2.14, P<0.05). When MTPα is knocked down in adult MB neurons, the effect of associative massed training on ATP consumption was lost (n=9-10, t_17_=0.07, P=0.94). **f-h.** Constitutive expression of the LD-GFP transgene in MB neurons revealed the presence of LDs in the somas of MB neurons (indicated by arrowheads, white box inset: magnified area, maximum z-stack projection of 3 confocal slices; total z-axis: 1 µm; scale bars=15 µm, inset scale bar=4 µm). Genetically impairing FA mitochondrial import (CPT1 KD) or β-oxidation (MTPα KD) in MB neurons increased LD density in somas of MB neurons (f: n=7-8, F_2,19_=26.51, P<0.0001), whereas the volume of individual LDs did not change (g: =4-8, F_2,16_=0.71, P=0.051). Data are expressed as the mean ± s.e.m. with dots as individual values, and analyzed by one-way analysis of variance (ANOVA) with post hoc testing by Newman–Keuls pairwise comparisons test (a-c, f-g) or by unpaired two-sided t-test (d). Asterisks refer to the least significant P value of post hoc comparison between the genotype of interest and the genotypic controls (a- c), or to the P value of post hoc comparison between the genotype of interest and the genotypic control (f-g), or to the P value of the unpaired t-test comparison (d). *P<0.05, ***P < 0.001, ****P<0.0001, NS: not significant.

Once inside the mitochondria, activated FAs can be oxidized into acetyl-CoA via the β-oxidation pathway, a cyclic process that generates reduced NADH and FADH_2_ and requires the successive action of different chain length-specific enzymes for the complete oxidation of FAs^19,22^ (Fig. 1a). We therefore investigated the role of two major enzymes in this complex pathway: the mitochondrial trifunctional protein (MTP), which shortens long-chain FAs to medium-chain lengths^22,23^; and HAD1, an enzyme more specific to short-chain FAs, with homology to mammalian 3-hydroxyacyl-CoA dehydrogenases^22,24^. Downregulation of either MTPα (the α subunit of MTP) or HAD1 expression in adult MB neurons caused a specific memory defect after massed training (Fig. 1c-d, Extended Data Fig. 1c-f, Supplementary Tables 1-2). These results show that the β-oxidation pathway is required in MB neurons to sustain memory formation after massed training.

Since FA β-oxidation is associated with ATP production^22^, we investigated whether ATP production in MB neurons was increased in flies forming memory after massed training as compared to control flies subjected to a non-associative massed protocol in which the presentations of electric shocks and odor are dissociated in time (unpaired protocol). To image ATP levels *in vivo*, we expressed in MB neurons the genetically-encoded ATP FRET sensor AT1.03NL, a modified version of the widely used ATeam ATP sensor that is optimized for use in non-homeotherm species such as Drosophila^25^. Following sodium azide application, which blocks the mitochondrial respiratory chain, the FRET ratio of the ATP sensor showed a rapid drop, the slope of which can be used to estimate the level of ATP consumption (Extended Data Fig. 1g). Notably, sodium azide application had no effect on the FRET signal of a nonfunctional version of the probe^25^ that is insensitive to ATP level (Extended Data Fig. 1g). After associative massed training, the rate of sodium azide-induced ATP decrease was higher than that observed in flies subjected to the non-associative massed protocol, indicating that associative massed training increases the consumption of ATP produced by mitochondria in the soma of MB neurons (Fig. 1e). When MTPα was knocked down in adult MB neurons, the effect of associative massed training on ATP consumption was lost (Fig. 1e), showing that FA β-oxidation sustains the increased ATP production in soma of MB neurons triggered by associative massed training.

To further support our finding that FA β-oxidation occurs *in vivo* in MB neurons, we investigated whether the impaired import of FAs to mitochondria and their subsequent oxidation results in FA accumulation. Because free FAs in the cytoplasm are toxic^26^, they are stored as triacylglycerol (TAG) in lipid droplets^27^. In order to reveal the accumulation of FA, we used the LD-GFP transgene consisting of a GFP fused to the lipid droplet (LD) domain of Klar, a protein involved in LD transport^10,28^. In flies constitutively expressing the LD marker LD-GFP under the control of a specific MB neuronal driver, only a few LDs were detected in the soma of MB neurons, similar to what was previously described^10^. In contrast, genetically impairing FA mitochondrial import (CPT1 KD) or β- oxidation (MTPα KD) in MB neurons led to the abnormal accumulation of LDs in their somas, with no significant effect on the individual size of LDs (Fig. 1f-h). These results show that impairing FAs import and oxidation by mitochondria in MB neurons result in their intracellular accumulation.

We have shown that FAs are imported into the mitochondria to be oxidized in order to support ATP production in neurons, a critical process for memory formed after massed training (Fig. 1). As shown in Fig. 1f-g, even if LDs are difficult to detect, they are nevertheless present in MB neurons of naive flies. If MB neuronal LD are a source of FAs for β-oxidation for memory after massed training, lipase activity should then be required. We therefore aimed to identify a lipase that would mediate the lipolysis of LD triacylglycerol into glycerol and FAs in MB neurons. In other cells, this process occurs through the sequential action of three FA lipases^27^. Interestingly, Brummer (Bmm), the orthologue of mammalian adipose triglyceride lipase (ATGL)^29^, which catalyzes the first hydrolysis reaction, is expressed in MB neurons (Extended Data Fig. 2a). When Bmm expression is knocked down in MB neurons, we observed LD accumulation in their somas (Fig. 2a-c). This result demonstrates that in MB neurons of naïve flies Bmm mediates LD lipolysis. Therefore, we hypothesize that memory formed after massed training requires the activity of this lipase to deliver FAs from neuronal LDs for mitochondrial oxidation. To test this hypothesis, we investigated memory formed after massed training in flies expressing a Bmm RNAi in adult MB neurons. Downregulation of Bmm expression in the adult MB induced a specific memory impairment after massed training (Fig. 2d, Extended Data Fig. 2b-c, Supplementary Table 3). These results indicate that Bmm-mediated lipolysis in MB neurons is specifically required to sustain memory formed after massed training.

**Figure 2:**
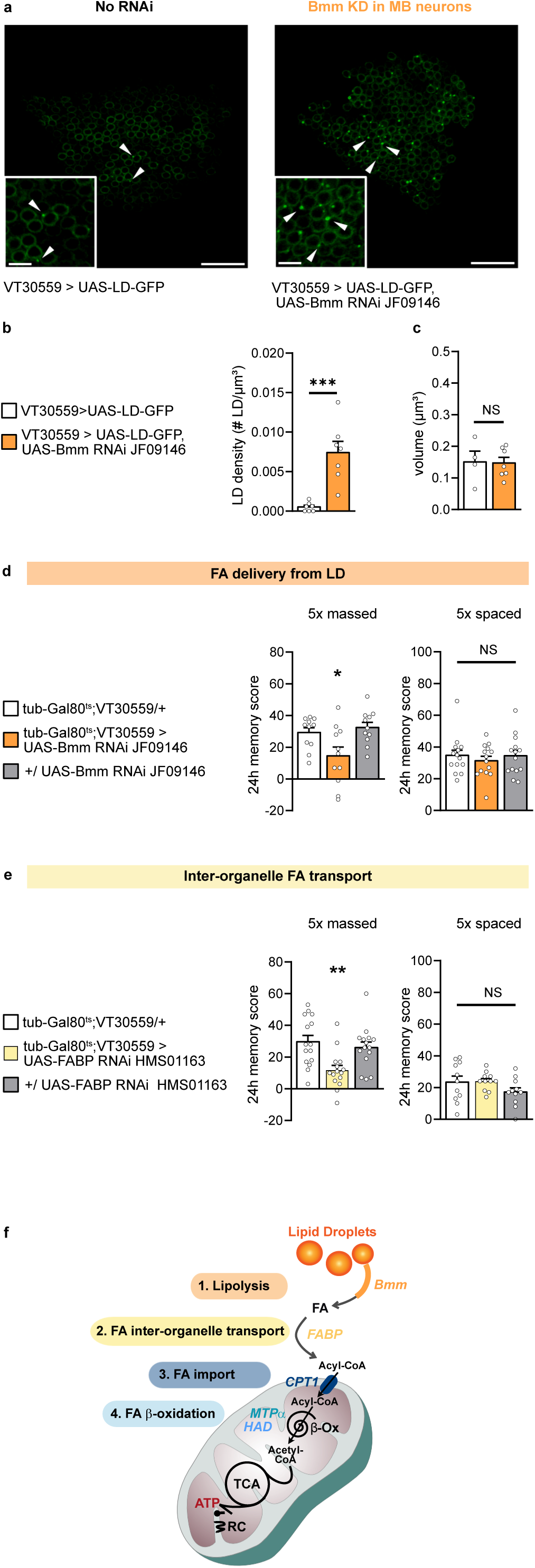
FAs derived from MB neuronal lipid stores are used to sustain memory formed after massed training. **a-c**. Genetically impairing LD lipolysis (Bmm KD) in MB neurons increased LD density in the somas of MB neurons compared to the genotypic control (b: n=7, t_12_=4.78, P<0.001), whereas the volume of individual LDs did not change (c: =4-7, t_9_=0.09, P=0.93). LDs are indicated by arrowheads, white box inset: magnified area, maximum z-stack projection of 3 confocal slices (total z-axis: 1µm), scale bars=15 µm, inset scale bar=4 µm. **d.** Bmm KD in adult MB neurons impaired memory after massed training (n=11, F_2,30_=5.33, P<0.05), but not after spaced training (n=14, F_2,39_=0.36, P=0.70). **e.** FABP KD in adult MB neurons impaired memory after massed training (n=15, F_2,42_=7.22, P<0.01), but not after spaced training (n=11, F_2,30_=1.74, P=0.19). **f.** Schema of the identified actors of FA metabolism required for memory formed after massed training. Data are expressed as the mean ± s.e.m. with dots as individual values, and analyzed by one-way analysis of variance (ANOVA) with post hoc testing by Newman–Keuls pairwise comparisons test (d-e) or by unpaired two-sided t-test (b-c). Asterisks refer to the least significant P value of post hoc comparison between the genotype of interest and the genotypic controls (d- e) or to the P value of the unpaired t-test comparison (b-c). *P<0.05, **P < 0.01, ***P < 0.001, NS: not significant.

To facilitate their transport between organelles and limit free FA toxicity, FAs rely on FA-binding proteins (FABPs), which are particularly critical for FA transport from LDs to mitochondria^30,31^. We therefore investigated whether this LD-to-mitochondria transport is also critical for memory formed after massed training. Whereas several FABPs exist in mammals, only one orthologue is found in Drosophila^32^. We observed that FABP downregulation in adult MB neurons caused a specific memory defect after massed training (Fig. 2e, Extended Data Fig. 2d-e, Supplementary Table 2, 3). These results show that FA inter-organelle transport mediated by FABP is required in neurons to support memory formed after massed training. Based on these data, we propose a neuronal pathway of FA use, from neuronal LD stores to mitochondrial energy production, which is employed specifically during memory formation following cramming-like learning (Fig. 2f).

Interestingly, it has been shown *in vitro* that under specific conditions such as starvation, in which fibroblasts become more reliant on FA β-oxidation, mitochondria are remodeled into highly connected networks, and these elongated mitochondria are required for appropriate FA flux to mitochondria^30^. Thus, we asked whether mitochondria network remodeling is also involved to support memory formed after massed training. We first investigated whether associative massed training triggers mitochondrial network remodeling. Specifically, we used high-resolution 3D-STED microscopy to image mitochondria of flies expressing in MB neurons a DsRed addressed to the mitochondrial matrix, as in one of our previous work^33^. We characterized mitochondria volume distribution in the soma of MB neurons of brains fixed 1 h after massed training (see Methods for details). These mitochondria were categorized into four groups based on their volume (see Extended Data Fig. 3a for the distribution of mitochondria volume used to define the limit of these categories), and the density of mitochondria in each category (number of mitochondria per µm^3^) was compared between experiments. When flies were subjected to associative massed training, the density of the largest mitochondria increased in MB somas as compared to control flies subjected to the non-associative massed protocol (Fig. 3a), while the density of intermediate mitochondria remained unchanged and the density of the smallest mitochondria was reduced. These data demonstrate that associative massed training triggers mitochondrial network remodeling in the MB soma. Three non-exclusive mechanisms could result in the observed changes: an increase in fusion of small mitochondria into larger mitochondria; an increase in mitochondria biogenesis; and the increased transport of small mitochondria to other neuronal compartments. We previously showed that impairing mitochondria microtubule-mediated transport does not affect memory formed after massed training, whereas it is required for memory formed after spaced training^33^. Thus, even if we cannot exclude that some mitochondria transport occurs from the soma to other compartments, it is probably not the main mitochondria network remodeling process triggered by massed training to support memory formation. On the other hand, mitochondrial biogenesis is a complex process involving gene expression and membrane synthesis, which can take several hours to a few days^34,35^. The duration of this process is therefore hardly compatible with the timeframe of our experiments in which we observed an increase in large mitochondria in the MB soma of flies already one hour after a 20-min learning session. Altogether, these data suggest that the increase in large mitochondria in the MB soma triggered by associative massed training relies mainly on the mitochondria fusion process.

**Figure 3:**
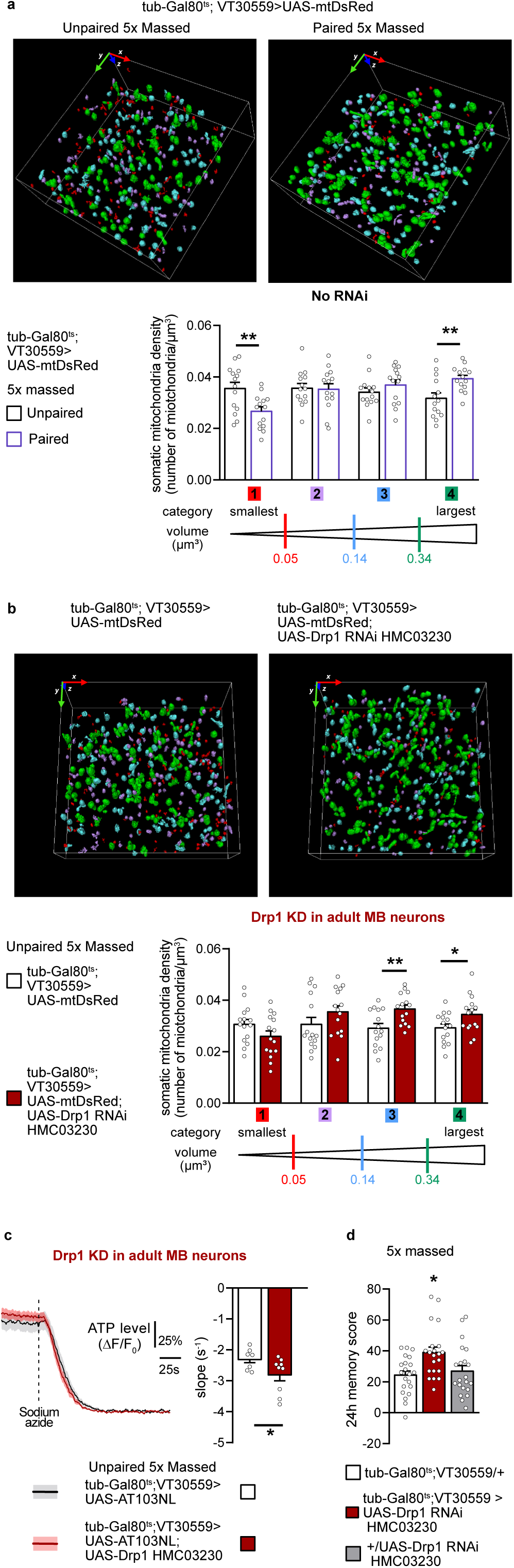
Mitochondrial network remodeling facilitates ATP production in the soma of MB neurons and improves memory after massed training. **a.** Mitochondria were categorized according to their volume into 4 groups. Detected mitochondria are shown with their color-coded categories (smallest mitochondria: category 1, red; small intermediate: category 2, purple; medium intermediate: category 3, blue; largest mitochondria: category 4, green). In order to compare between the different conditions, we calculated the density of mitochondria for each category by normalizing the number of mitochondria in each category by the volume of the minimal envelope containing all of the detected mitochondria in the ROI (see Methods for details). One hour after training, in flies subjected to 5x massed associative training, the density of the largest mitochondria was increased in MB somas compared to control flies subjected to the non-associative massed protocol (largest mitochondria category (4): n=14, t_26_=3.08, P<0.01). The intermediate categories (2 and 3) were not significantly affected by associative massed training (intermediate mitochondria category (2): n=14, t_26_=0.15, P=0.88; (3): n=14, t_26_=1.12, P=0.27), whereas the density of the smallest mitochondria was decreased upon associative training (smallest mitochondria category (1): n=14, t_26_=3.01, P<0.01.). **b.** When Drp1 was knocked down in adult MB, the density of large mitochondria in MB neuronal somas was increased as compared to the genotypic control (category 3: n=15, t_28_=3.11, P<0.01; category 4: n=15, t_28_=2.17, P<0.05), whereas the density of small mitochondria did not change (category 1: n=15, t_28_=1.64, P=0.11; category 2: n=15, t_28_=1.34, P=0.19). **c.** When Drp1 was knocked down in adult MB neurons, the level of ATP consumption was increased as compared to the genotypic control (n=9, t_16_=2.28, p<0.01). **d.** Drp1 KD in adult MB neurons increased memory performance after massed training (n=21, F_2,60_=5.55, P<0.01). Data are expressed as the mean ± s.e.m. with dots as individual values, and analyzed by one-way analysis of variance (ANOVA) with post hoc testing by Newman–Keuls pairwise comparisons test (d) or by an unpaired two-sided t-test (a-c). Asterisks refer to the least significant P value of post hoc comparison between the genotype of interest and the genotypic controls (d) or to the P value of the unpaired t-test comparison (a-c). *P<0.05, **P < 0.01, NS: not significant.

To further investigate the role of fusion in remodeling the mitochondria network in MB somas upon massed training, we shifted the mitochondria fission/fusion equilibrium toward fusion by downregulating the expression of the main actor of mitochondrial fission, Drp1 (Dynamin-related protein 1)^36^. First, we characterized the effect of Drp1 KD in adult MB neurons on mitochondria morphology in the soma of those neurons. Using the same approach as in Fig. 3a, we observed, in flies expressing Drp1 RNAi together with the mitochondrial DsRed in adult MB, that the density of large mitochondria is increased in the soma as compared to the genotypic controls (Fig. 3b). As elongated mitochondria have been associated with increased metabolic capacity, we investigated ATP production when Drp1 was knocked down in adult MB neurons. Using the same approach as in Fig. 1, we measured an increase in ATP consumption in the soma of MB neurons of Drp1 KD flies as compared to the genotypic control flies (Fig. 3c). These results suggest that ATP production is elevated in the MB neuronal soma of Drp1 KD flies. We then investigated whether this genetically-induced increase in mitochondria size in MB neuronal somas associated with the increase in ATP level affects memory formed after massed training. Strikingly, downregulating Drp1 expression in the adult MB resulted in an improved memory performance after massed training (Fig. 3d). In contrast, memory formed after spaced training was impaired in those flies (Extended Data Fig. 3b). In the absence of RNAi expression, memory formed after either massed training or spaced training was normal (Extended Data Fig. 3b). Finally, sensory perceptions were normal in induced flies (Supplementary Table 4). These results were replicated with a second non-overlapping Drp1 RNAi line (Extended Data Fig. 3c, Supplementary Table 4). Because increased mitochondria size has been shown to decrease mitochondria motility^37,38^ and mitochondria axonal transport in MB neurons is required to support memory formed after spaced training^33^, the observed impairment in memory formed after spaced training in Drp1 KD flies is likely related to decreased mitochondria motility in MB neurons. More importantly, our data establish that the favoring of mitochondrial metabolic capacity (here by knocking down Drp1 in MB neurons) improves memory performance at the behavioral level, after massed training. Thus, while the loss-of-function experiments presented earlier (Figs. 1 and 2) could be interpreted as supportive of neuronal activity within a classical framework of cellular energy metabolism, this result additionally shows that boosting energy production *per se* can be sufficient to improve cognitive performance.

Our study challenges the long-standing view that glucose and its byproducts are the main energy providers for neurons in the healthy brain^26,39,40^, revealing that FAs are used *in vivo* by neurons as an energy source to support memory formation. However, the mitochondrial oxidation of FAs can also be accompanied by a concomitant increase in ROS production^41^. Deciphering whether neurons that use FAs for energy purposes have a specific ROS defense system, or if using FAs has a detrimental cost, will need further investigation.

We identified the triacylglycerol lipase Bmm, the orthologue of mammalian ATGL, as a critical step in LD usage to support memory formation. Previous studies have focused on the role of DDHD2, a lipase exhibiting both di- and tri-acylglycerol hydrolase activities that act together with ATGL in LD lipolysis^42,43^, and have shown that it is required *in vitro* for ATP production in neurons^11,12^. DDHD2 deficiency in humans is associated with cognitive impairment^44^, and in line with this, a constitutive KO mouse model exhibits long-term memory deficit^45^. However, because FA metabolism is critical for other aspects of neuronal physiology such as membrane turnover and protein acylation in order to regulate gene expression, it was difficult to conclude from these studies if FAs generated from LD lipolysis are used for energy production in neurons. Here, we demonstrated using a neuron-specific KD that *in vivo* LD lipolysis in neurons generates FAs that are used to produce energy by mitochondrial oxidation during memory formation. Whether the FAs stored in neuronal LDs originate from the surrounding glia or from peripheral organs (such as the fat body) in flies is an open question. Recently, it was shown in the Drosophila larva brain that lipid transport from astrocytes to neurons is mediated by astrocyte-derived apolipoproteins coupled to the neuronal receptor LpR1^46^. Determining if the same actors are at play in the adult brain to provide FAs for energy production to support memory formation will require further investigation.

Altogether, our data demonstrate that neuronal FA oxidation is required in MB neurons to sustain the formation of memory induced by cramming-like learning, thus uncovering a metabolic flexibility in neurons to support memory formation. In addition to providing new perspectives in the brain energy metabolism field, the improvement in memory performance after cramming-like learning by increasing mitochondria metabolic capacity provides strong support for the concept of mitochondrial plasticity as a key factor of memory robustness^47^.

## Methods

### Fly strains

*Drosophila melanogaster* flies were raised on standard food medium containing yeast, cornmeal and agar, on a 12 h:12 h light-dark cycle at 18°C and 60% humidity. The Canton Special (CS) strain was used as the wild-type strain. All lines were outcrossed for at least three generations to flies carrying a CS wild-type background. Pan-neuronal expression of transgenes was achieved using the *elav-Gal4* line^48^. For transgene expression in MB neurons, we used the *VT30559-*Gal4 (VDRC: v206077) line^14^. In order to restrict UAS/GAL4-mediated expression to the adult stage, we used the TARGET system^21^ with the *tubulin-GAL80^ts^* (BDSC: 7019) line to construct the inducible driver line *tubulin-GAL80^ts^; VT30559-Gal4*, previously described in ^14^. Gal4 activity was released by transferring 1–2-day-old adult flies to 30°C for 2-3 days. In some behavioral experiments (Extended Data Figs. 1d and 2b), the UAS-Dicer2 transgene (BDSC: 24650) was used in combination with the MB inducible driver (Tub-Gal80ts; UAS-Dcr2, VT30559-Gal4) to increase either MTPα RNAi GD11299 or Bmm RNAi GD5139 efficiency, an approach that our laboratory successfully used in a previous study^17^.

The following UAS-transgene lines were obtained from the Vienna *Drosophila* Resource Center (VDRC): *UAS-Bmm RNAi GD5139* (VDRC: v37877)^10^, *UAS-FABP RNAi KK116001* (VDRC: v109169), *UAS-CPT1 RNAi KK100935* (VDRC: v105400)^49^, *UAS-MTPα* RNAi *GD11299* (VDRC: v21845)^50^, and *UAS-Drp1 GD10456* (VDRC: v44155). The following UAS-transgene lines were obtained from the Bloomington *Drosophila* Stock Center (BDSC): *UAS-Bmm RNAi JF01946* (BDSC:25926)^10^, *UAS-FABP RNAi HMS01163 (BDSC: 34685)*^51^*, UAS-CPT1 RNAi HMS00040* (BDSC:34066)^49^, *UAS-RNAi MTPα HMS00660* (BDSC: 32873)^52^, *UAS-RNAi HAD1 HMC05280* (BDSC: 62273), *Had1^nl^* (BDSC: 1037)^24^, and *UAS-Drp1 HMC03230* (BDSC: 51483). Reporter lines used in this study include: *UAS- AT1.03NL* and *UAS-AT1.03RK*^25^, *UAS-mtDsRed* (BDSC: 93056)^53^, *Mi{Trojan-GAL4.1}bmm^MI13321-TG4.1^* (BDSC:67510), and *UAS-mcD8::GFP* ^54^. The *UAS-LD-GFP* line was a kind gift from M. A. Welte^28^, and *UAS-AT1.03NL* and *UAS- AT1.03RK* were provided by H. Imamura^25^. The following lines were generated in this study: *UAS-LD-GFP, UAS-Bmm RNAi JF01946* and *UAS-LD-GFP, UAS-CPT1 RNAi HMS00040* and *UAS-LD-GFP, UAS-MTPα RNAi HMS00660* and *UAS-AT1.03NL; UAS-MTPα RNAi HMS00660* and *UAS-AT1.03NL, UAS-Drp1 HMC03230* and *UAS-mtDsRed, UAS-Drp1 HMC03230*. For RNAi lines not yet validated, the efficiency of each RNAi construct to decrease the mRNA level of the targeted gene was confirmed following the protocol detailed in the Quantitative PCR analyses section (see below), with results presented in Supplementary Table 2.

### Olfactory conditioning and memory test

The behavior experiments, including sample sizes, were conducted similarly to previous studies from our research group^14,15,33^. For RNAi induction, 1–2-day-old flies were kept at 30°C for 2-3 days until conditioning. The non-induced control flies, in which RNAi expression is inhibited, were kept at 18°C. For all experiments, training and testing were performed at 25°C and 80% humidity; after conditioning, flies were kept at 18°C until testing. Briefly, groups of approximately 30-40 flies were subjected to one of the following olfactory conditioning protocols: five consecutive associative training cycles (5x massed), or five associative cycles spaced by 15-min inter-trial intervals (5x spaced). Custom-built barrels allowing parallel training of up to 6 groups were used for conditioning. Throughout the conditioning protocol, each barrel was plugged into a constant airflow at 2 L·min^-1^. The sequence of one conditioning cycle consisted of an initial 90-s period of non-odorized airflow, followed by 60 s of the conditioned odor paired with 12 pulses of electric shocks (60V; 1 pulse every 5 s, pulse duration: 1.2 s). After 45 s of non-odorized airflow, the second odor was presented for 60 s without electric shocks, followed by 45 s of non-odorized airflow. The odorants 3-octanol (>95% purity; Fluka 74878, Sigma-Aldrich) and 4- methylcyclohexanol (99% purity; Fluka 66360) were diluted in paraffin oil at 0.360 mM and 0.325 mM, respectively, and were alternately used as conditioned stimuli. During unpaired conditionings, the odor and shock stimuli were delivered separately in time, with shocks starting 3 min before the first odorant.

The memory test was performed in a T-maze apparatus, typically after 24 h of massed or spaced training. Flies were exposed simultaneously to both odorants for 1 min in the dark. The performance index (PI) was calculated as the number of flies attracted to the unconditioned odor minus the number of flies attracted to the conditioned odor, divided by the total number of flies in the experiment, and the resulting number was multiplied by 100. A single PI value is the average of two scores obtained from two groups of genotypically identical flies conditioned in two reciprocal experiments, using either odorant (3-octanol or 4- methylcyclohexanol) as the conditioning stimulus. The indicated “n” is the number of independent PI values for each genotype.

Olfactory avoidance and shock avoidance tests were conducted similarly to previous studies from our research group^14,15,33^. The shock response tests were performed at 25°C by placing flies in two connected compartments; electric shocks were administered in only one of the compartments. Flies were given 1 min to move freely in these compartments, after which they were trapped, collected, and counted. The compartment in which the electric shocks were delivered was alternated between two consecutive groups. Shock avoidance was calculated as for the memory test. Since the delivery of electric shocks can modify olfactory acuity, our olfactory avoidance tests were performed on flies that had first been presented another odor paired with electric shocks. Innate odor avoidance was measured in a T-maze similar to those used for memory tests, in which one arm of the T-maze was connected to a bottle containing odor diluted in paraffin oil, and the other arm was connected to a bottle with paraffin oil only. Naive flies were given the choice between the two arms during 1 min. The odor-interlaced side was alternated for successively tested groups. Odor concentrations used in this assay were the same as for the memory assays. At these concentrations, both odorants are innately repulsive.

### *In vivo* ATP imaging

As in all previous imaging studies from our lab^14,15,33^, *in vivo* imaging experiments were performed in female flies due to their larger size, which facilitates surgery. Briefly, female flies carrying a *TubGal80^ts^;VT30559-GAL4* construct were crossed to CS males or to males carrying either the *UAS-AT1.03NL* transgene, or the *UAS- AT1.03RK* transgene or the appropriate UAS-RNAi together with the *UAS-AT1.03NL* transgene. Crosses for imaging experiments were raised at 23°C to avoid RNAi expression during development. The 1–2-day-old adult progeny were induced for 3 days at 30°C. After either the associative or non-associative massed protocol, a single fly was affixed to a plastic coverslip using a nontoxic dental glue (Protemp II 3M ESPE), and 90 μL of an artificial hemolymph solution were added on top of the coverslip. The composition of the artificial hemolymph solution was: 130 mM NaCl (Sigma, S9625), 5 mM KCl (Sigma, P3911), 2 mM MgCl_2_ (Sigma, M9272), 2 mM CaCl_2_ (Sigma, C3881), 5 mM D-trehalose (Sigma, T9531), 30 mM sucrose (Sigma, S9378), and 5 mM HEPES-hemisodium salt (Sigma, H7637). Surgery was performed as previously described^14,15,33^ to expose the brain for optical imaging. At the end of the surgery, a fresh 90-μL drop of the appropriate saline solution was applied on the aperture in the fly’s head. Two-photon imaging was performed on a Leica TCS-SP5 upright microscope equipped with a 25x, 0.95 NA water immersion objective. Two-photon excitation of mTFP was achieved using a Mai Tai DeepSee laser tuned to 820 nm. 512 × 200 images were acquired at a frame rate of two images per second, and the entire duration of each recording was 360 s. After 2 min of baseline acquisition, 10 µL of a 50 mM sodium azide solution (Sigma cat. #71289; prepared in the same artificial hemolymph solution) were injected into the 90-mL droplet bathing the fly’s brain, bringing sodium azide to a final concentration of 5 mM. To analyze the ATP imaging experiments, ROIs were delimited by hand around each visible MB soma region, and the average intensity of the mTFP and Venus channels over each ROI was calculated over time after background subtraction. The ATP sensor is designed so that FRET from CFP to YFP increases as the ATP concentration increases, and thus the FRET ratio was calculated as YFP intensity divided by CFP intensity. This ratio was normalized by a baseline value calculated over 120 s, starting at 120 s after the drug injection (corresponding to the last 120 s of the recording), when the ATP level was below the detection limit of the sensor. The slope was calculated between 90% and 30% of the plateau, which allowed the level of ATP consumption to be estimated, using the same principle as in ^14^ to measure MB pyruvate consumption. Imaging analysis was performed using a custom-written MATLAB script^14^. The indicated ‘n’ is the number of animals assayed in each condition.

### Immunostaining, image acquisition and analysis for STED microscopy

The immunostaining for STED experiments as well as the image acquisitions and analysis were conducted similarly to our previous study^33^. Flies were raised at 18°C; to induce RNAi expression as well as mt-DsRed, adult flies were kept at 30.5°C for 3 d before conditioning with either the 5x massed associative or non-associative protocol. One hour after the end of the training period, whole adult flies were fixed in 4% paraformaldehyde (Electron Microscopy Sciences) in PBST (PBS containing 0.6% Triton X-100) at 4°C overnight. Next, brains were dissected in PBS solution and fixed again for 1 h at room temperature (RT) in 4% paraformaldehyde in PBST followed by three 20-min rinses in PBST, with blocking for 2 h at RT with 2% BSA (Sigma-Aldrich cat#A9085) in PBST. Samples were incubated with the primary antibody Atto647N FluoTag®-X4 anti-RFP (NanoTag cat#N0404- Atto647N-L) at 1:100 in the blocking buffer at 4°C overnight. The next day, brains were rinsed twice for 20 min in PBST and once for 20 min in PBS. Samples were then mounted in ProLong Gold Antifade Mountant (Invitrogen) using precision cover glasses (thickness: No. 1.5H; Marienfeld Superior).

Single-color 3D-STED imaging was performed using a Leica TCS SP8 STED 3X microscope (excitation at 633 nm, STED depletion laser at 775 nm (pulsed), with 60% in x and y dimensions and 50% in z dimension) with a 93x motCorr glycerol immersion objective (NA=1.3). Images of one MB soma region per brain were obtained with a voxel size of 41.68 nm (x) x 41.68 nm (y) x 72.48 nm (z). One ROI per brain containing only somas was delimited from the full image (120 µm x 120 µm x 8 µm) using a constant square box of 21 µm x 21 µm x 8 µm. The raw ROIs were then analyzed using Icy Software and the HK-mean plugin followed by the connected-component (ROI extraction mode, removal of the border objects) plugin to detect mitochondria and their volumes^55,56^. Each ROI typically contained 200-350 mitochondria. The same parameters (*i.e.* the Gaussian pre-filter and the intensity classes) were used for all ROIs analyzed in the study. These parameters were first determined on a set of ROIs from brain samples by a researcher blind for the genotype and the behavioral training. The volume of the minimal envelope containing all of the detected mitochondria in each ROI was determined using a custom-written ImageJ macro^57^. Briefly, we proceeded with the binarization of single fluorescent images of the z-stack series obtained from the Icy analysis. Binarized fluorescent objects were then connected using XOR and Convex Hull mathematical operators. Surface and volume estimates were performed using the 3D object option available in Fiji.

Based on the mitochondria volume distribution in the control group (5x massed unpaired, tub-Gal80^ts^>UAS- mtDsRed), we defined 4 categories of mitochondria, from the smallest volume to the largest, using the mean of the 25^th^ and 75^th^ percentiles and the medians of mitochondria volumes in each ROI from one fly brain (Extended Data Fig. 3a). The limit of each category is defined as follows: for category 1 (the smallest), the upper limit is the mean of the 25^th^ percentiles; for category 2, the lower limit is the mean of the 25^th^ percentiles and the upper limit is the mean of the medians; for category 3, the lower limit is the mean of the medians and the upper limit is the mean of the 75^th^ percentiles; and for category 4 (the largest), the lower limit is the mean of the 75^th^ percentiles. The same volume limits were used to define the categories in all experimental conditions. To allow comparisons between the samples (for each ROI from one fly brain), we normalized the number of mitochondria in each category by the volume of the minimal envelope containing all of the detected mitochondria, in order to obtain the mitochondria density for each category.

### LD staining and image analysis

To achieve LD staining in MB neurons, VT30559-GAL4 flies were crossed with flies bearing UAS-LD-GFP with or without the appropriate UAS-RNAi, and raised at 25°C. 1–3-day-old whole adult flies were fixed in 4% paraformaldehyde (Electron Microscopy Sciences) in PBST (PBS containing 1% Triton X-100) at 4°C overnight. Next, brains were dissected in PBS solution and fixed again for 1 h at room temperature (RT) in 4% paraformaldehyde in PBST. Brains were then rinsed once in PBST for 20 min, and twice in PBS for 20 min. After rinsing, brains were mounted using Prolong Mounting Medium (Invitrogen). Images of one MB soma region per brain were acquired the following day with a voxel size of 0.117 µm (x) x 0.117 µm (y) x 0.41 µm (z) on an Olympus FV1000 confocal microscope with a 60x/1.35 oil immersion objective using a 473 nm laser. Confocal z-stacks were imported into Fiji^58^ and CellProfiler 4.2.6 software^59^ for further analyses. Using Fiji, one ROI per brain containing only MB somas was delimited from the full image using a constant rectangular box of 120 µm x 90 µm x 2 µm. Next, CellProfiler was used for 3D object detection and counting to identify LDs from the background staining of the plasma membrane. Thresholding was automatically performed based on the Robust background thresholding method. This method assumes that the background distribution approximates a Gaussian distribution as we previously carried out for the LD analysis^17^. Specifically, the threshold background parameters used for the lower and upper outlier fractions were fixed to 0.02 and 0.01 respectively, with a mean averaging method, and a variance method with the standard deviation set to 4 and a threshold correction factor of 5. The watershed module was used to detect the 3D objects, and a filter was applied based on the object mean intensity to discard residual background from non-specific labeling of the plasma membrane (minimum intensity value: 0.025). The number of LDs and their volume were obtained using the measure object intensity and measure object size modules. To allow comparisons between the samples, we normalized the number of LDs by the volume of the MB soma region. This volume was estimated based on the background signal from the plasma membrane. Here, the object is defined using a manual thresholding that sets the lower limit to 0.00005, image smothering with a sigma of 1, and applying a median filter (window of 10) and then the “removeholes” module, with the “measure image area occupied” module to measure the volume occupied by the MB soma region. The indicated ‘n’ is the number of animals that were assayed in each condition.

### Immunohistochemistry experiments

Female flies carrying the *Mi{Trojan-GAL4.1}bmm^MI13321-TG4.1^*construct^60^ were crossed to male flies carrying the *UAS-mCD8::GFP* construct^54^. Prior to dissection, 2–4-day-old female flies were fixed in 4% paraformaldehyde in PBST (PBS containing 1% of Triton X-100) at 4°C overnight. Fly brains were dissected on ice in PBS solution and rinsed three times for 20 min in PBST. Then, brains were blocked with 2% BSA in PBST for 2 h. Next, samples were incubated with primary antibodies in the blocking solution (2% BSA in PBST) at 4°C overnight. The following primary antibodies were used: 1:250 rabbit anti-GFP (Invitrogen, A11122), and 1:100 mouse anti-nc82 (DSHB, nc82). The following day, brains were rinsed 3 times for 20 min with PBST and then incubated for 3 h at room temperature with secondary antibodies diluted in blocking solution. The following secondary antibodies were used: 1:400 anti-rabbit conjugated to Alexa Fluor 488 (Invitrogen, A11034), and 1:400 anti-mouse conjugated to Alexa Fluor 594 (Invitrogen, A11005). Brains were then rinsed once in PBST for 20 min, and twice in PBS for 20 min. After rinsing, brains were mounted using Prolong Mounting Medium (Invitrogen). Acquisitions were made with a Nikon A1R confocal microscope with a 40x/1.15 water immersion objective.

### Quantitative PCR analyses

To assess the efficiency of each RNAi line used in this study, quantitative PCR analyses were conducted similarly to previous studies from our research group^17,33^. Female flies carrying the *elav-Gal4* pan-neuronal driver were crossed with males carrying the specified UAS-RNAi or with CS males. Fly progeny were reared at 25°C throughout their development. Then, 1–2-day-old flies were transferred to fresh food for 1 day prior to RNA extraction. RNA extraction, DNAse I treatment (when needed) and cDNA synthesis were performed as previously^17^ using the RNeasy Plant Mini Kit (Qiagen), RNA MiniElute Cleanup Kit (Qiagen), DNAse I treatment (BioLabs), oligo(dT)20 primers and the SuperScript III First-Strand kit (Thermo Fisher Invitrogen). Amplification was performed using a LightCycler 480 (Roche) and the SYBR Green I Master mix (Roche). Specific primers used for each gene cDNA and the reference α-Tub84B (Tub, CG1913) cDNA are provided in Supplementary Table 5. Reactions were performed in triplicate. The specificity and size of amplification products were assessed by melting curve analyses. The level of cDNA for each gene of interest was compared against the level of the α- Tub84B reference cDNA. Expression relative to the reference was expressed as a ratio (2^-ΔCp^, where Cp is the crossing point).

### Statistical analysis

Data are expressed as the mean ± s.e.m. with dots as individual values (one experimental replicate, n=1) corresponding to: a group of 40-50 flies analyzed together in a behavioral assay; the response of a single recorded fly for ATP imaging; one brain for LD experiments or for mitochondria morphology analysis; or one mRNA extraction from heads of a group of 50 flies used for an RT-qPCR experiment. Statistical analysis was performed using the GraphPad Prism 8.0 software (GraphPad Software, La Jolla, CA, USA). Comparisons between two groups were performed by unpaired two-sided Student’s *t*-test, with results given as the value t*_x_* of the t distribution, where *x* is the number of degrees of freedom. Comparisons among three groups were performed by one-way analysis of variance (ANOVA) with post hoc testing by the Newman-Keuls pairwise comparisons test between the experimental group and its controls (significance: p<0.05). ANOVA results are given as the value of the Fisher distribution F_(*x,y*)_, where *x* is the number of degrees of freedom numerator and *y* is the total number of degrees of freedom denominator. Asterisks in each figure refer to the least significant post hoc comparison between the genotype of interest and the genotypic controls. The nomenclature used corresponds to *P<0.05, **P<0.01, ***P< 0.001, ****P<0.0001, ns: not significant, P>0.05. Figures were made using Affinity Designer v2.

## Data availability

No datasets that require mandatory deposition into a public database were generated during the current study. Processed data from imaging experiments and raw data of behavioral assays, will be provided as Source data file upon publication. Unprocessed images, which represent a large volume, will be available through e-mailing the corresponding authors, and will be shared without restriction.

## Code availability

No code was generated during this study.

## Acknowledgments

We thank the TRiP consortium at Harvard Medical School for providing transgenic RNAi fly stocks. We thank M. A. Welte for providing the UAS-LD-GFP strain and H. Imamura for the UAS-AT1.03NL and AT1.03RK strains. We thank Alexandre Didelet and Christelle Beauchamp for technical support with fly food preparation, Julia Minatchy for assistance with behavioral experiments, Amandine Delecourt for assistance with lipid droplet imaging experiments, and Lucie Cagninacci for assistance with RT-qPCR experiments. This work was funded by the European Research Council (ERC Advanced Grant EnergyMemo n°741550, to T.P.), and by the Agence Nationale de la Recherche to A.P. (ANR-22-CE16-0020) and P.-Y.P (ANR-20-CE92-0047-01). B.S. was funded by a doctoral fellowship from the École des Neurosciences de Paris (ENP) and a postdoctoral fellowship from Labex MemoLife.

## Author contributions

Conceptualization: A.P., T.P., B.S., and P.Y.P.; investigation: A.P. and B.S.; methodology: A.P., P.Y.P., T.P., B.S., and L.D.; writing/original draft preparation: A.P. and B.S.; writing/review and editing: A.P., B.S., T.P., and P.Y.P.; supervision: A.P., T.P., and P.Y.P.; funding acquisition: T.P., P.Y.P and A.P.

---

**Extended Data Figure 1:**
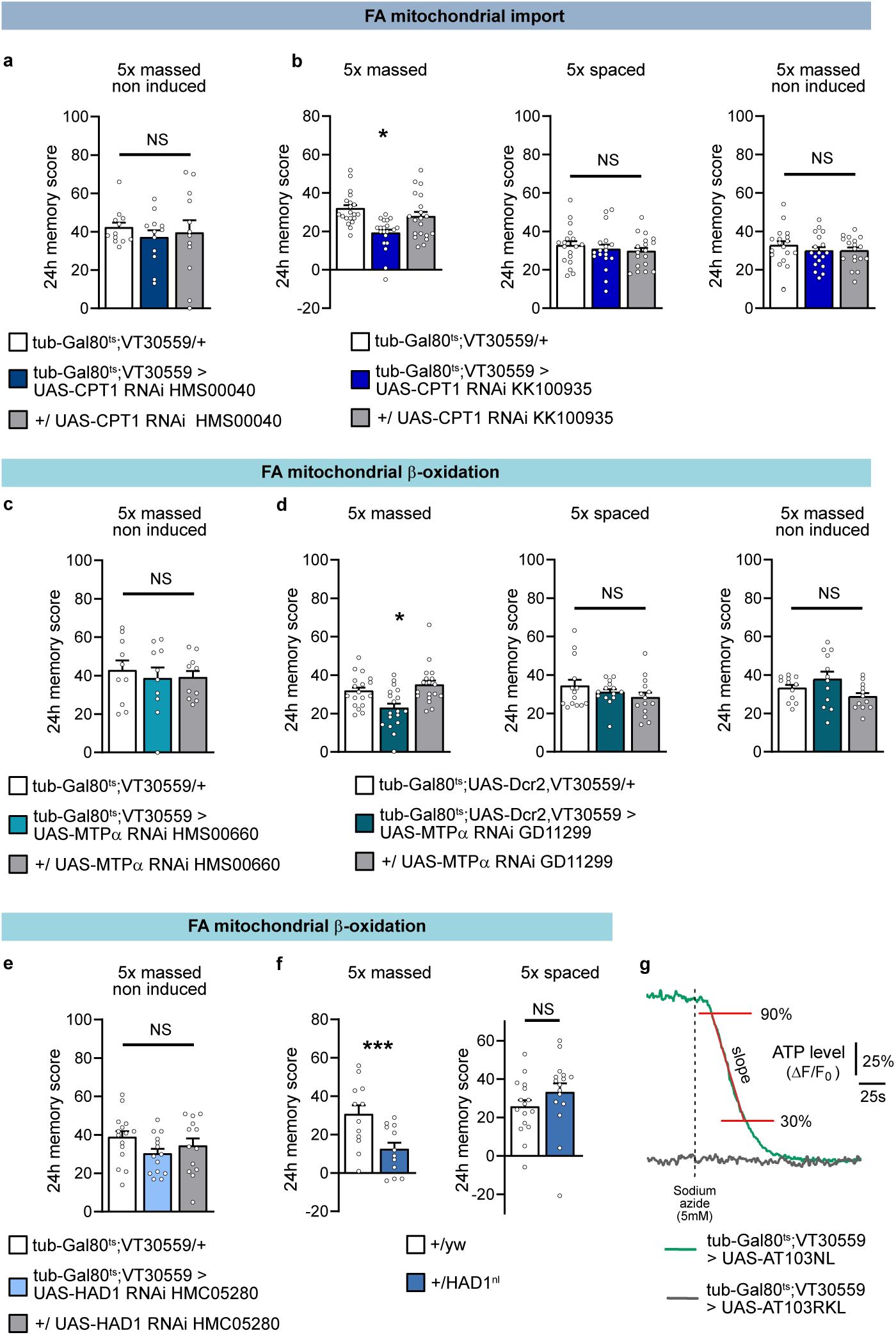
Control experiments for FA mitochondrial import and β-oxidation in MB neurons to sustain memory formed after massed training. **a.** When CPT1 RNAi expression was not induced, memory after massed training was normal (n=12, F_2,33_=0.30, P=0.74). **b**. A second non-overlapping RNAi targeting CPT1 (CPT1 RNAi KK10935) was used to confirm the specific defect in memory after massed training. Inhibition of CPT1 expression in adult MB neurons using CPT1 RNAi KK10935 impaired memory after massed training (n=20, F_2,57_=8.91, P<0.001), whereas memory after spaced training was normal (n=18, F_2,51_=0.43, P=0.66). Non-induced flies showed no memory defect after massed training (n=18, F_2,51_=0.61, P=0.55). **c.** When MTPα RNAi expression was not induced, memory after massed training was normal (n=10, F_2,27_=0.21, P=0.82). **d.** A second non-overlapping RNAi targeting MTPα (MTPα RNAi GD11299) was used to confirm the specific defect in memory after massed training. In order to increase RNAi efficiency, Dicer2 expression was induced together with the RNAi expression in adult MB neurons using the Tub-Gal80ts; UAS-Dcr2, VT30559 line. Inhibition of MTPα expression in adult MB neurons using this RNAi impaired memory after massed training (n=18, F_2,51_=7.30, P<0.001), whereas memory after spaced training was normal (n=14, F_2,39_=1.21, P=0.31). Non-induced flies showed no memory defect after massed training (n=12, F_2,33_=2.62, P=0.09). **e.** When HAD1 RNAi expression was not induced, memory after massed training was normal (n=13-15, F_2,39_=1.63, P=0.21). **f.** We used a HAD1 null mutant to confirm HAD1 function in memory formed after massed training. Since HAD1^nl^ is in a yw background, we used heterozygous +/yw flies as a control for the +/HAD1^nl^ experimental group. Heterozygous +/HAD1^nl^ flies had a memory defect after massed training as compared to control +/yw flies (n=12, t_22_=2.97, P<0.01), whereas memory after spaced training was normal (n=16, t_30_=1.23, P=0.23). **g.** Applying 5 mM of sodium azide (dashed line) resulted in a fast decrease in the FRET ratio of the ATP FRET sensor AT1.03NL expressed in adult MB neurons, whereas the FRET ratio of AT1.03RKL did not change (mean trace ± s.e.m.). The slope of the FRET ratio was measured between 90% and 30% of the baseline level (plateau reached after sodium azide application corresponding to the lower limit of the FRET sensor detection). Data are expressed as the mean ± s.e.m. with dots as individual values, and analyzed by one-way analysis of variance (ANOVA) with post hoc testing by Newman–Keuls pairwise comparisons test (a-e) or by unpaired two-sided t-test (f-g). Asterisks refer to the least significant P value of post hoc comparison between the genotype of interest and the genotypic controls (a-e), or to the P value of the unpaired t-test comparison (f-g). *P<0.05, ***P < 0.001, NS: not significant.

**Extended Data Figure 2:**
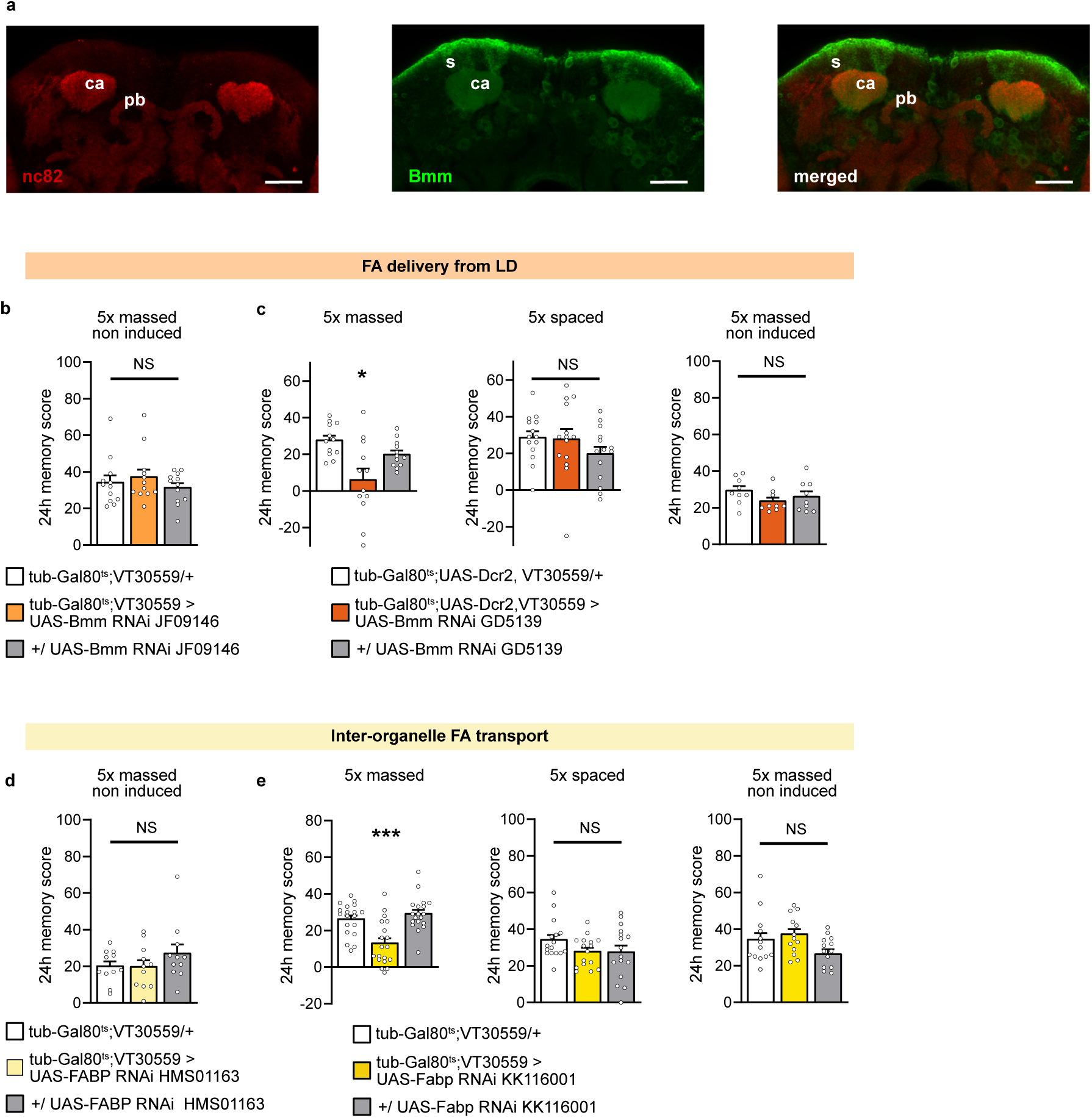
Control experiments for FA delivery from LDs and transport to mitochondria in MB neurons to sustain memory formed after massed training. **a.** Immunohistochemistry of *Mi{Trojan-GAL4.1}bmm^MI13321-TG4.1^*>UAS-mCD8::GFP brains showing the Bmm expression pattern in green (GFP) and pan-neuronal anti-nc82 counterstaining in red. Images are a maximum intensity projection of 3 confocal planes (total z-axis: 2 µm) of the soma and calyx MB region of the Drosophila brain (posterior brain). Clear Bmm expression can be detected in the soma and the calyx of MB neurons (s: soma of MB neurons, Ca: Calyx of MB, pb: protocerebral bridge of the central complex). Scale bar: 30 µm. **b.** When Bmm RNAi expression was not induced, memory after massed training was normal (n=12, F_2,33_=0.67, P=0.52). **c.** A second non-overlapping RNAi targeting Bmm (Bmm RNAi GD5139) was used to confirm the specific defect in memory after massed training. As previously carried out in ^17^ using this RNAi line, Dicer2 expression was induced together with RNAi expression in adult MB neurons (Tub-Gal80ts; UAS-Dcr2, VT30559 line) to increase RNAi efficiency. Inhibition of Bmm expression in adult MB neurons using this RNAi impaired memory after massed training (n=12, F_2,33_=7.21, P<0.01), whereas memory after spaced training was normal (n=14, F_2,39_=1.24, P=0.30). Non-induced flies showed no memory defect after massed training (n=9, F_2,24_=1.50, P=0.24). **d.** When FABP RNAi expression was not induced, memory after massed training was normal (n=11, F_2,30_=1.15, P=0.33). **e.** A second non-overlapping RNAi targeting FABP (FABP RNAi KK116001) was used to confirm the specific defect in memory after massed training. Inhibition of FABP expression in adult MB neurons using this RNAi impaired memory after massed training (n=19, F_2,54_=13.51, P<0.0001), whereas memory after spaced training was normal (n=16, F_2,45_=1.72, P=0.19). Non-induced flies showed no memory defect after massed training (n=13, F_2,36_=2.12, P=0.13). Data are expressed as the mean ± s.e.m. with dots as individual values, and analyzed by one-way analysis of variance (ANOVA) with post hoc testing by Newman–Keuls pairwise comparisons test (b-e). Asterisks refer to the least significant P value of post hoc comparison between the genotype of interest and the genotypic controls (b-e). *P<0.05, ***P < 0.001, NS: not significant.

**Extended Data Figure 3:**
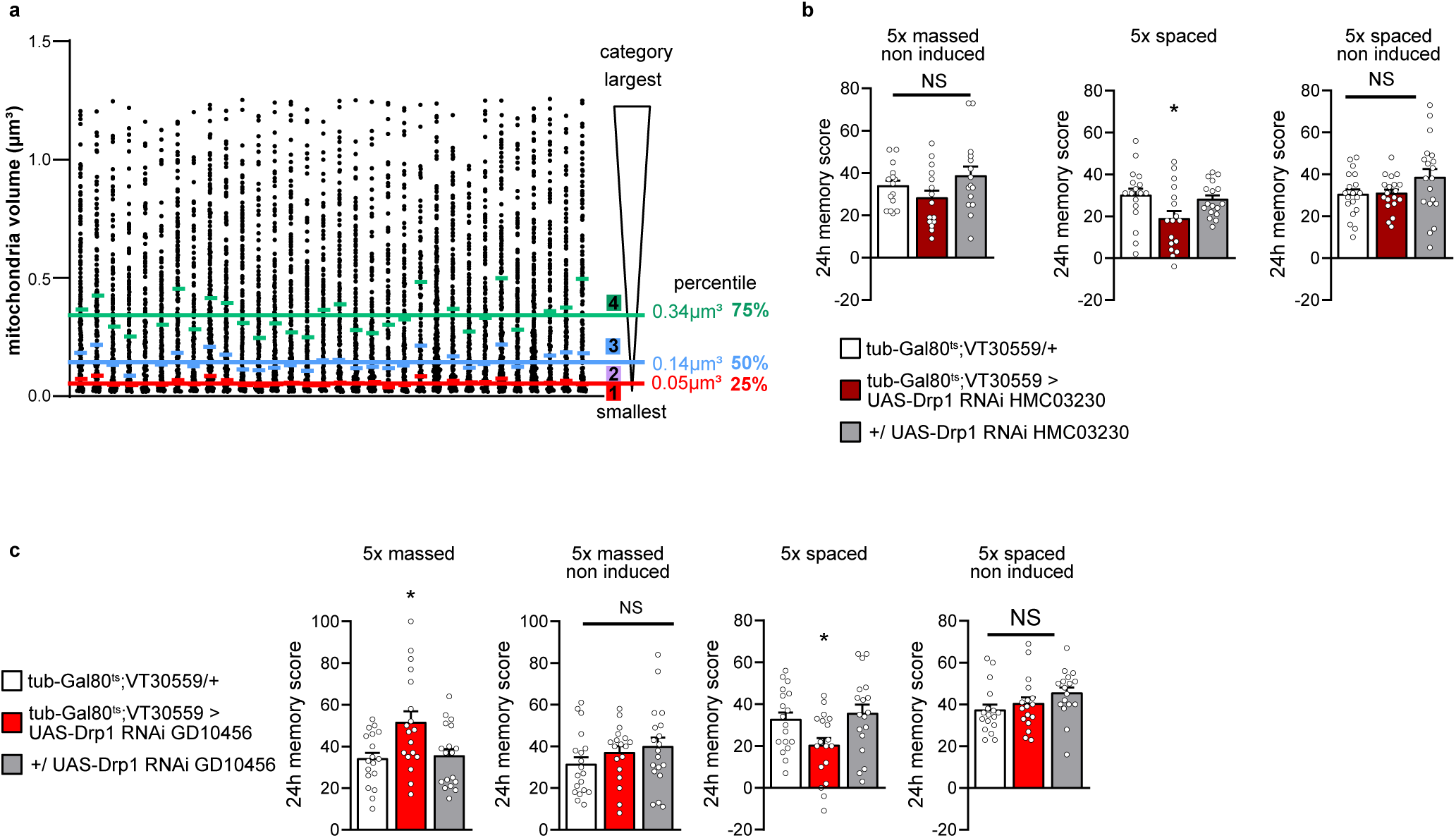
Control experiments for mitochondrial network remodeling facilitates ATP production in the soma of MB neurons and improves memory after massed training. **a.** The mitochondria volume distribution is shown for each ROI of all unpaired control flies used in this study, where each dot represents the volume of one single identified mitochondrion. The means of the quartile limits from all control flies subjected to the massed unpaired protocol (*i.e.* tub-Gal80^ts^; VT30559>UAS-mtDsRed flies subjected to the unpaired 5x massed protocol in Fig. 3a-b) were used to define the 4 mitochondria categories. **b.** When Drp1 RNAi expression was not induced, memory after massed training was normal (n=15, F_2,42_=2.01, P=0.15). Drp1 KD in adult MB neurons impaired memory after spaced training (n=17, F_2,48_=3.68, P<0.05). When Drp1 RNAi expression was not induced, memory after spaced training was normal (n=19, F_2,54_=2.25, P=0.12). **c.** A second non-overlapping RNAi targeting Drp1 (Drp1 RNAi GD10456) was used to confirm the specific increase in memory after massed training. Inhibition of Drp1 expression in adult MB neurons using this RNAi increased memory performance after massed training (n=18, F_2,51_=5.53, P<0.01), whereas memory after spaced training was impaired (n=18, F_2,51_=4.38, P<0.05). Non-induced flies showed normal memory performance after either massed training (n=18-19, F_2,53_=1.33, P=0.27) or spaced training (n=17, F_2,48_=2.01, P=0.15).

## References

1. Beattie, D. S. & Basford, R. E. Brain Mitochondria—Iii Fatty Acid Oxidation by Bovine Brain Mitochondria. Journal of Neurochemistry 12, 103–111 (1965).

2. Yang, S. Y., He, X. Y. & Schulz, H. Fatty acid oxidation in rat brain is limited by the low activity of 3-ketoacyl-coenzyme A thiolase. Journal of Biological Chemistry 262, 13027–13032 (1987).

3. Volk, M. E., Millington, R. H. & Weinhouse, Sidney. OXIDATION OF ENDOGENOUS FATTY ACIDS OF RAT TISSUES IN VITRO. Journal of Biological Chemistry 195, 493–501 (1952).

4. Ebert, D., Haller, R. G. & Walton, M. E. Energy Contribution of Octanoate to Intact Rat Brain Metabolism Measured by ^13^ C Nuclear Magnetic Resonance Spectroscopy. J. Neurosci. 23, 5928– 5935 (2003).

5. Edmond, J., Robbins, R. A., Bergstrom, J. D., Cole, R. A. & De Vellis, J. Capacity for substrate utilization in oxidative metabolism by neurons, astrocytes, and oligodendrocytes from developing brain in primary culture. J. Neurosci. Res. 18, 551–561 (1987).

6. Ioannou, M. S. et al. Neuron-Astrocyte Metabolic Coupling Protects against Activity-Induced Fatty Acid Toxicity. Cell 177, 1522–1535.e14 (2019).

7. Morant-Ferrando, B. et al. Fatty acid oxidation organizes mitochondrial supercomplexes to sustain astrocytic ROS and cognition. Nat Metab 5, 1290–1302 (2023).

8. C, F., et al. Cell-type-specific profiling of brain mitochondria reveals functional and molecular diversity. Nature neuroscience 22, (2019).

9. Schönfeld, P. & Reiser, G. Brain energy metabolism spurns fatty acids as fuel due to their inherent mitotoxicity and potential capacity to unleash neurodegeneration. Neurochemistry International 109, 68–77 (2017).

10. Wat, L. W. et al. A role for triglyceride lipase brummer in the regulation of sex differences in Drosophila fat storage and breakdown. PLoS Biol 18, e3000595 (2020).

11. Kumar, M., et al. DDHD2 Is Necessary for Activity-Driven Fatty Acid Fueling of Nerve Terminal Function. http://biorxiv.org/lookup/doi/10.1101/2023.12.18.572201 (2023) doi:10.1101/2023.12.18.572201.

12. Saber, S. H. et al. DDHD2 provides a critical flux of saturated fatty acids to support neuronal energy demands. preprint (2024) doi:: 10.1101/2023.12.31.573799.

13. Tully, T., Preat, T., Boynton, S. C. & Del Vecchio, M. Genetic dissection of consolidated memory in Drosophila. Cell 79, 35–47 (1994).

14. Plaçais, P.-Y. et al. Upregulated energy metabolism in the Drosophila mushroom body is the trigger for long-term memory. Nat Commun 8, 15510 (2017).

15. Comyn, T., Preat, T., Pavlowsky, A. & Plaçais, P.-Y. PKCδ is an activator of neuronal mitochondrial metabolism that mediates the spacing effect on memory consolidation. eLife 13, (2024).

16. Heisenberg, M. Mushroom body memoir: from maps to models. Nat Rev Neurosci 4, 266–275 (2003).

17. Silva, B. et al. Glia fuel neurons with locally synthesized ketone bodies to sustain memory under starvation. Nat Metab 4, 213–224 (2022).

18. McGarry, J. D. & Brown, N. F. The Mitochondrial Carnitine Palmitoyltransferase System — From Concept to Molecular Analysis. European Journal of Biochemistry 244, 1–14 (1997).

19. Carillo, M. R. et al. L-Carnitine in Drosophila: A Review. Antioxidants 9, 1310 (2020).

20. McGarry, J. D., Leatherman, G. F. & Foster, D. W. Carnitine palmitoyltransferase I. The site of inhibition of hepatic fatty acid oxidation by malonyl-CoA. Journal of Biological Chemistry 253, 4128–4136 (1978).

21. McGuire, S. E. Spatiotemporal Rescue of Memory Dysfunction in Drosophila. Science 302, 1765– 1768 (2003).

22. Houten, S. M., Violante, S., Ventura, F. V. & Wanders, R. J. A. The Biochemistry and Physiology of Mitochondrial Fatty Acid β-Oxidation and Its Genetic Disorders. Annu. Rev. Physiol. 78, 23–44 (2016).

23. Kishita, Y., Tsuda, M. & Aigaki, T. Impaired fatty acid oxidation in a *Drosophila* model of mitochondrial trifunctional protein (MTP) deficiency. Biochemical and Biophysical Research Communications 419, 344–349 (2012).

24. Tobler, J. E. & Grell, E. H. Genetics and physiological expression of β-hydroxy acid dehydrogenase in Drosophila. Biochem Genet 16, 333–342 (1978).

25. Tsuyama, T. et al. In Vivo Fluorescent Adenosine 5′-Triphosphate (ATP) Imaging of Drosophila melanogaster and Caenorhabditis elegans by Using a Genetically Encoded Fluorescent ATP Biosensor Optimized for Low Temperatures. Anal. Chem. 85, 7889–7896 (2013).

26. Schönfeld, P. & Reiser, G. How the brain fights fatty acids’ toxicity. Neurochemistry International 148, 105050 (2021).

27. Olzmann, J. A. & Carvalho, P. Dynamics and functions of lipid droplets. Nature Reviews Molecular Cell Biology 20, 137–155 (2019).

28. Yu, Y. V., Li, Z., Rizzo, N. P., Einstein, J. & Welte, M. A. Targeting the motor regulator Klar to lipid droplets. BMC Cell Biology 12, 9 (2011).

29. Grönke, S. et al. Brummer lipase is an evolutionary conserved fat storage regulator in Drosophila. Cell Metabolism 1, 323–330 (2005).

30. Rambold, A. S., Cohen, S. & Lippincott-Schwartz, J. Fatty Acid Trafficking in Starved Cells: Regulation by Lipid Droplet Lipolysis, Autophagy, and Mitochondrial Fusion Dynamics. Developmental Cell 32, 678–692 (2015).

31. Furuhashi, M. & Hotamisligil, G. S. Fatty acid-binding proteins: role in metabolic diseases and potential as drug targets. Nat Rev Drug Discov 7, 489–503 (2008).

32. Gerstner, J. R., Vanderheyden, W. M., Shaw, P. J., Landry, C. F. & Yin, J. C. P. Fatty-Acid Binding Proteins Modulate Sleep and Enhance Long-Term Memory Consolidation in Drosophila. PLoS ONE 6, e15890 (2011).

33. Pavlowsky, A. et al. Spaced training activates Miro/Milton-dependent mitochondrial dynamics in neuronal axons to sustain long-term memory. Current Biology 34, 1904–1917.e6 (2024).

34. Cardanho-Ramos, C. & Morais, V. A. Mitochondrial Biogenesis in Neurons: How and Where. International Journal of Molecular Sciences 22, 13059 (2021).

35. Hodneland Nilsson, L. I., et al. A new live-cell reporter strategy to simultaneously monitor mitochondrial biogenesis and morphology. Sci Rep 5, 17217 (2015).

36. Westermann, B. Mitochondrial dynamics in model organisms: What yeasts, worms and flies have taught us about fusion and fission of mitochondria. Seminars in Cell & Developmental Biology 21, 542–549 (2010).

37. Berthet, A. et al. Loss of Mitochondrial Fission Depletes Axonal Mitochondria in Midbrain Dopamine Neurons. J. Neurosci. 34, 14304–14317 (2014).

38. Verstreken, P. et al. Synaptic Mitochondria Are Critical for Mobilization of Reserve Pool Vesicles at Drosophila Neuromuscular Junctions. Neuron 47, 365–378 (2005).

39. Schönfeld, P. & Reiser, G. Why does Brain Metabolism not Favor Burning of Fatty Acids to Provide Energy? - Reflections on Disadvantages of the Use of Free Fatty Acids as Fuel for Brain. J Cereb Blood Flow Metab 33, 1493–1499 (2013).

40. Panov, A., Orynbayeva, Z., Vavilin, V. & Lyakhovich, V. Fatty Acids in Energy Metabolism of the Central Nervous System. BioMed Research International 2014, 1–22 (2014).

41. Seifert, E. L., Estey, C., Xuan, J. Y. & Harper, M.-E. Electron transport chain-dependent and - independent mechanisms of mitochondrial H2O2 emission during long-chain fatty acid oxidation. J Biol Chem 285, 5748–5758 (2010).

42. Yu, L. Cooperation of acylglycerol hydrolases in neuronal lipolysis. Journal of Lipid Research 64, 100462 (2023).

43. Hofer, P. et al. Cooperative lipolytic control of neuronal triacylglycerol by spastic paraplegia-associated enzyme DDHD2 and ATGL. Journal of Lipid Research 64, 100457 (2023).

44. Schuurs-Hoeijmakers, J. et al. Mutations in DDHD2, encoding an intracellular phospholipase A(1), cause a recessive form of complex hereditary spastic paraplegia. American journal of human genetics 91, (2012).

45. Akefe, I. O. et al. The DDHD2-STXBP1 interaction mediates long-term memory via generation of saturated free fatty acids. The EMBO Journal 43, 533–567 (2024).

46. Yin, J. et al. Brain-specific lipoprotein receptors interact with astrocyte derived apolipoprotein and mediate neuron-glia lipid shuttling. Nat Commun 12, 2408 (2021).

47. Comyn, T., Preat, T., Pavlowsky, A. & Plaçais, P.-Y. Mitochondrial plasticity: An emergent concept in neuronal plasticity and memory. Neurobiology of Disease 203, 106740 (2024).

48. Luo, L., Liao, Y. J., Jan, L. Y. & Jan, Y. N. Distinct morphogenetic functions of similar small GTPases: Drosophila Drac1 is involved in axonal outgrowth and myoblast fusion. Genes Dev. 8, 1787–1802 (1994).

49. Tiwari, S. K., Toshniwal, A. G., Mandal, S. & Mandal, L. Fatty acid β-oxidation is required for the differentiation of larval hematopoietic progenitors in Drosophila. eLife 9, e53247 (2020).

50. McMullen, E. et al. Glycolytically impaired Drosophila glial cells fuel neural metabolism via β- oxidation. Nat Commun 14, 2996 (2023).

51. Kis, V., Barti, B., Lippai, M. & Sass, M. Specialized Cortex Glial Cells Accumulate Lipid Droplets in Drosophila melanogaster. PLoS ONE 10, e0131250 (2015).

52. Zipper, L., Batchu, S., Kaya, N. H., Antonello, Z. A. & Reiff, T. The MicroRNA miR-277 Controls Physiology and Pathology of the Adult Drosophila Midgut by Regulating the Expression of Fatty Acid β-Oxidation-Related Genes in Intestinal Stem Cells. Metabolites 12, 315 (2022).

53. Lutas, A., Wahlmark, C. J., Acharjee, S. & Kawasaki, F. Genetic Analysis in Drosophila Reveals a Role for the Mitochondrial Protein P32 in Synaptic Transmission. G3 Genes|Genomes|Genetics 2, 59–69 (2012).

54. Scheunemann, L., Plaçais, P.-Y., Dromard, Y., Schwärzel, M. & Preat, T. Dunce Phosphodiesterase Acts as a Checkpoint for Drosophila Long-Term Memory in a Pair of Serotonergic Neurons. Neuron 98, 350–365.e5 (2018).

55. de Chaumont, F. et al. Icy: an open bioimage informatics platform for extended reproducible research. Nat Methods 9, 690–696 (2012).

56. Dufour, A., Meas-Yedid, V., Grassart, A. & Olivo-Marin, J.-C. Automated quantification of cell endocytosis using active contours and wavelets. in 2008 19th International Conference on Pattern Recognition 1–4 (2008). doi:10.1109/ICPR.2008.4761748.

57. Bun, P. Minimal Enveloppe Estimation. Zenodo 10.5281/ZENODO.7118602 (2022).

58. Schindelin, J., et al. Fiji: an open-source platform for biological-image analysis. Nature Methods 9, 676–682 (2012).

59. Stirling, D. R. et al. CellProfiler 4: improvements in speed, utility and usability. BMC Bioinformatics 22, 433 (2021).

60. Lee, P.-T., et al. A gene-specific T2A-GAL4 library for Drosophila | eLife. 7, e35574 (2018).

