## Supplementary information for "Neuronal fatty acid oxidation fuels memory after intensive learning"

**Supplementary Tables: Control experiments for olfactory acuity and electric shock avoidance.**

Expression of the different RNAi constructs used in this study in MB neurons did not have any significant effect on olfactory acuity or the avoidance of electric shocks. The *P*-value indicated for RNAi-expressing flies is the lowest one obtained from the two pairwise comparisons between these flies and their driver (*tubulin-GAL80ts; VT30559/+*) or effector (*UAS-…RNAi/+*) controls. n is indicated for each experiment.

**Supplementary Table 1: Control experiments for olfactory acuity and electric shock avoidance related to Fig. 1 and Extended Data Fig. 1.**

| Genotype | Shock avoidance | | Naive odor avoidance | | | |
|---|---|---|---|---|---|---|
| | | | Octanol | | Methylcyclohexanol | |
| | Mean ± s.e.m. | Statistics | Mean ± s.e.m. | Statistics | Mean ± s.e.m. | Statistics |
| *TubGal80$^{ts}$;VT30559/+* | 0.55±0.10 | $F_{2,21}$=0.51 p=0.61 n=8 | 0.71±0.04 | $F_{2,21}$=0.027 p=0.97 n=8 | 0.55±0.02 | $F_{2,21}$=1.04 p=0.37 n=8 |
| *TubGal80$^{ts}$;VT30559> UAS-CPT1 HMS0040* | 0.64±0.11 | | 0.69±0.06 | | 0.68±0.08 | |
| *UAS-CPT1 HMS0040/+* | 0.49±0.10 | | 0.69±0.07 | | 0.67±0.08 | |
| *TubGal80$^{ts}$;VT30559/+* | 0.70±0.04 | $F_{2,45}$=0.35 p=0.71 n=16 | 0.49±0.05 | $F_{2,33}$=0.74 p=0.49 n=12 | 0.47±0.06 | $F_{2,33}$=0.14 p=0.87 n=12 |
| *TubGal80$^{ts}$;VT30559> UAS-CPT1 KK100935* | 0.72±0.04 | | 0.46±0.05 | | 0.50±0.06 | |
| *UAS-CPT1 KK100935/+* | 0.67±0.04 | | 0.42±0.03 | | 0.47±0.03 | |
| *TubGal80$^{ts}$;VT30559/+* | 0.43±0.07 | $F_{2,21}$=0.53 p=0.60 n=8 | 0.41±0.07 | $F_{2,21}$=0.28 p=0.76 n=8 | 0.40±0.08 | $F_{2,21}$=0.11 p=0.89 n=8 |
| *TubGal80$^{ts}$; VT30559> UAS-MTPα HMS00660* | 0.36±0.13 | | 0.35±0.07 | | 0.35±0.5 | |
| *UAS-MTPα HMS00660/+* | 0.30±0.07 | | 0.35±0.06 | | 0.38±0.09 | |
| *TubGal80$^{ts}$; UAS-Dcr2, VT30559/+* | 0.70±0.06 | $F_{2,27}$=0.36 p=0.70 n=10 | 0.56±0.04 | $F_{2,39}$=1.78 p=0.18 n=14 | 0.57±0.05 | $F_{2,39}$=0.21 p=0.81 n=14 |
| *TubGal80$^{ts}$; UAS-Dcr2, VT30559> UAS-MTPα GD11299* | 0.66±0.04 | | 0.50±0.03 | | 0.53±0.04 | |
| *UAS-MTPα GD11299/+* | 0.71±0.05 | | 0.48±0.03 | | 0.53±0.05 | |
| *TubGal80$^{ts}$;VT30559/+* | 0.51±0.04 | $F_{2,27}$=0.28 p=0.76 n=10 | 0.60±0.11 | $F_{2,24}$=0.86 p=0.44 n=9 | 0.77±0.09 | $F_{2,24}$=0.18 p=0.84 n=9 |
| *TubGal80$^{ts}$; VT30559> UAS-HAD1 HMC05280* | 0.51±0.10 | | 0.40±0.11 | | 0.70±0.08 | |

| | | | | | | |
|---|---|---|---|---|---|---|
| *UAS-HAD1 HMC05280/+* | 0.57±0.06 | | 0.45±0.10 | | 0.76±0.10 | |
| *+/yw* | 0.41±0.11 | $t_{14}$=0.78 p=0.45 n=8 | 0.62±0.07 | $t_{14}$=0.41 p=0.69 n=8 | 0.54±0.11 | $t_{14}$=1.60 p=0.13 n=8 |
| *+/HAD1*$^{nl}$ | 0.51±0.08 | | 0.66±0.06 | | 0.75±0.07 | |

**Supplementary Table 2: Efficiency of genetic knockdowns used in the study.**

Statistical comparisons were made using a two-sided unpaired t-test. Asterisks illustrate the significance level, with the following nomenclature: ***P<0.001, **P<0.01; *P<0.05.

| Genotype | Mean ± s.e.m. | Statistics | % of mRNA reduction |
|---|---|---|---|
| *elav/+* | 1.05±0.07 | $t_4$=7.461 p=0.0017 (**) n=3 | 79% |
| *elav>UAS-CPT1 HMS0040* | 0.23±0.09 | | |
| *elav/+* | 0.77±0.67 | $t_4$=4.150 p=0.0060 (**) n=4 | 60% |
| *elav>UAS-CPT1 KK100935* | 0.31±0.09 | | |
| *elav/+* | 0.90±0.27 | $t_6$=2.707 p=0.0353 (*) n=4 | 80% |
| *elav>UAS-MTPα HMS00660* | 0.18±0.01 | | |
| *elav/+* | 1.02±0.26 | $t_4$=3.256 p=0.0312 (*) n=3 | 84% |
| *elav>UAS-MTPα GD11299* | 0.16±0.03 | | |
| *elav/+* | 1.09±0.07 | $t_4$=5.626 p=0.0049 (**) n=3 | 75% |
| *elav>UAS-HAD1 HMC05280* | 0.27±0.12 | | |
| *+/yw* | 1.19±0.12 | $t_4$=3.578 p=0.0232 (*) n=3 | 50% |
| *+/$Had1^{nl}$* | 0.59±0.11 | | |
| *elav/+* | 2.09±0.25 | $t_6$=4.365 p=0.047 (**) n=4 | 57% |
| *elav>UAS-FABP HMS01163* | 0.89±0.10 | | |
| *elav/+* | 2.33±0.21 | $t_6$=7.597 p=0.0003 (***) n=4 | 75% |
| *elav>UAS-FABP KK116001* | 0.59±0.10 | | |
| *elav/+* | 0.08±0.01 | $t_5$=3.281 p=0.0219 (*) n=3-4 | 38% |
| *elav>UAS-Drp1 HMC03230* | 0.05±0.00 | | |
| *elav/+* | 0.08±0.01 | $t_5$=2.699 p=0.0428 (*) n=3-4 | 38% |
| *elav>UAS-Drp1 GD10456* | 0.05±0.01 | | |

**Supplementary Table 3: Control experiments for olfactory acuity and electric shock avoidance related to Fig. 2 and Extended Data Fig. 2.**

| Genotype | Shock avoidance | | Naive odor avoidance | | | |
|---|---|---|---|---|---|---|
| | | | Octanol | | Methylcyclohexanol | |
| | Mean ± s.e.m. | Statistics | Mean ± s.e.m. | Statistics | Mean ± s.e.m. | Statistics |
| *TubGal80$^{ts}$;VT30559/+* | 0.52±0.05 | $F_{2,45}$=1.70 p=0.19 n=16 | 0.51±0.06 | $F_{2,39}$=1.41 p=0.26 n=14 | 0.39±0.03 | $F_{2,39}$=1.37 p=0.27 n=14 |
| *TubGal80$^{ts}$;VT30559> UAS-Bmm JF09146* | 0.62±0.05 | | 0.65±0.07 | | 0.47±0.05 | |
| *UAS-Bmm JF09146/+* | 0.50±0.04 | | 0.57±0.06 | | 0.37±0.04 | |
| *TubGal80$^{ts}$; UAS-Dcr2, VT30559/+* | 0.37±0.06 | $F_{2,33}$=0.11 p=0.90 n=12 | 0.76±0.04 | $F_{2,33}$=1.9 p=0.17 n=12 | 0.73±0.05 | $F_{2,29}$=1.22 p=0.31 n=10-11 |
| *TubGal80$^{ts}$; UAS-Dcr2, VT30559> UAS-Bmm GD5139* | 0.39±0.05 | | 0.65±0.05 | | 0.64±0.05 | |
| *UAS-Bmm GD5139/+* | 0.42±0.10 | | 0.74±0.05 | | 0.74±0.04 | |
| *TubGal80$^{ts}$;VT30559/+* | 0.35±0.07 | $F_{2,21}$=1.06 p=0.36 n=8 | 0.38±0.06 | $F_{2,21}$=0.08 p=0.92 n=8 | 0.32±0.10 | $F_{2,21}$=0.19 p=0.83 n=8 |
| *TubGal80$^{ts}$;VT30559> UAS-FABP HMS01163* | 0.53±0.13 | | 0.40±0.06 | | 0.35±0.09 | |
| *UAS-FABP HMS01163/+* | 0.48±0.06 | | 0.41±0.04 | | 0.39±0.06 | |
| *TubGal80$^{ts}$;VT30559/+* | 0.65±0.02 | $F_{2,21}$=0.44 p=0.65 n=8 | 0.47±0.06 | $F_{2,33}$=0.73 p=0.49 n=12 | 0.43±0.05 | $F_{2,33}$=0.19 p=0.83 n=12 |
| *TubGal80$^{ts}$;VT30559> UAS-FABP KK116001* | 0.62±0.05 | | 0.56±0.07 | | 0.45±0.05 | |
| *UAS-FABP KK116001/+* | 0.60±0.04 | | 0.47±0.5 | | 0.48±0.06 | |

**Supplementary Table 4: Control experiments for olfactory acuity and electric shock avoidance related to Fig. 3 and Extended Data Fig. 3.**

| Genotype | Shock avoidance | | Naive odor avoidance | | | |
|---|---|---|---|---|---|---|
| | | | Octanol | | Methylcyclohexanol | |
| | Mean ± s.e.m. | Statistics | Mean ± s.e.m. | Statistics | Mean ± s.e.m. | Statistics |
| *TubGal80$^{ts}$;VT30559/+* | 0.43±0.08 | $F_{2,45}$=0.61 p=0.55 n=16 | 0.57±0.06 | $F_{2,45}$=0.64 p=0.53 n=16 | 0.52±0.07 | $F_{2,45}$=0.94 p=0.40 n=16 |
| *TubGal80$^{ts}$;VT30559> UAS-Drp1 HMC03230* | 0.51±0.06 | | 0.61±0.05 | | 0.60±0.05 | |
| *UAS-Drp1 HMC03230/+* | 0.39±0.08 | | 0.66±0.05 | | 0.64±0.06 | |
| *TubGal80$^{ts}$;VT30559/+* | 0.41±0.06 | $F_{2,69}$=0.49 p=0.61 n=24 | 0.50±0.05 | $F_{2,2}$=0.19 p=0.83 n=21-22 | 0.59±0.07 | $F_{2,45}$=0.70 p=0.50 n=16 |
| *TubGal80$^{ts}$;VT30559> UAS-Drp1 GD10456* | 0.48±0.08 | | 0.46±0.06 | | 0.60±0.05 | |
| *UAS-Drp1 GD10456/+* | 0.50±0.06 | | 0.52±0.08 | | 0.68±0.05 | |

**Supplementary Table 5: Primer sequences used for quantitative PCR.**

| Gene | Related to Figure: | Forward primer | Reverse primer |
|---|---|---|---|
| *α-Tub84B* | Fig. 1 and Extended Data Fig. 1 | TTGTCGCGTGTGAAACACTTC | CTGGACACCAGCCTGACCAAC |
| *CPT1* | Fig. 1 and Extended Data Fig. 1 | ACCATTGAGTCATCCGCCTG | CATCCTGGTAGCCCAACTGG |
| *MTPα* | Fig. 1 and Extended Data Fig. 1 | CAGGAACCTCCAAGGACACC | TTCCTGCAGCAGTCTGATGG |
| *HAD1* | Fig. 1 and Extended Data Fig. 1<br>*HAD1 RNAi HMC05280* | AATGGGACCGACACCGAAAA | CTCAACTGGGTGAGGCAGTT |
| | Extended Data Fig. 1<br>*Mutant Had1nl* | CAGGTGGTGCTGTACGACAT | GAGATGCAGGCGAATTGCTG |
| *FABP* | Fig. 2 and Extended Data Fig. 2 | TCAACAGTGAACATCTTGTGC | GCTGTTGCCCATCTTGCG |
| *Drp1* | Fig. 3 and Extended Data Fig. 3 | ATCTACAGCCCACTCGATGAT | GAAGCACTTCTTGGTGTGCAG |
